# A speed limit on serial strain replacement from original antigenic sin

**DOI:** 10.1101/2024.01.04.574172

**Authors:** Lauren McGough, Sarah Cobey

## Abstract

Many pathogens evolve to escape immunity, yet it remains difficult to predict whether immune pressure will lead to diversification, serial replacement of one variant by another, or more complex patterns. Pathogen strain dynamics are mediated by cross-protective immunity, whereby exposure to one strain partially protects against infection by antigenically diverged strains. There is growing evidence that this protection is influenced by early exposures, a phenomenon referred to as original antigenic sin (OAS) or imprinting. In this paper, we derive new constraints on the emergence of the pattern of successive strain replacements demonstrated by influenza, SARS-CoV-2, seasonal coronaviruses, and other pathogens. We find that OAS implies that the limited diversity found with successive strain replacement can only be maintained if *R*_0_ is less than a threshold set by the characteristic antigenic distances for cross-protection and for the creation of new immune memory. This bound implies a “speed limit” on the evolution of new strains and a minimum variance of the distribution of infecting strains in antigenic space at any time. To carry out this analysis, we develop a theoretical model of pathogen evolution in antigenic space that implements OAS by decoupling the antigenic distances required for protection from infection and strain-specific memory creation. Our results demonstrate that OAS can play an integral role in the emergence of strain structure from host immune dynamics, preventing highly transmissible pathogens from maintaining serial strain replacement without diversification.

## Introduction

Antigenically variable pathogens consist of immunologically distinct strains whose spatiotemporal dynamics depend on the pathogen’s capacity to diversify and on the extent to which host immune responses elicited by one strain protect against infection with other strains. This interdependence gives rise to qualitatively distinct patterns of diversity or strain structures [1, 2, 3, 4, 5, 6]. Examples include pathogens with high antigenic diversity at the host population scale (e.g., *Neisseria meningitidis* [7, 8, 9, 10], enteroviruses [11]) and others with lower antigenic diversity at any given time but fast turnover (e.g., influenza A in humans [12, 13], seasonal coronaviruses [14], SARS-CoV-2 [15, 16, 17], and others [18]).

Successive strain replacement typifies the dynamics of many common respiratory pathogens, yet it only arises in transmission models in some conditions [12, 6, 19, 20, 1, 21, 2, 22, 23, 24, 12, 25, 26, 27, 28, 29, 30]. Large individual-based models have found that low antigenic diversity and fast turnover result from low mutation rates and strong cross-immunity between strains [31, 6]. Short-term strain-transcending immunity [12, 27] and punctuated antigenic changes [32, 19] have been hypothesized to be essential for serial antigenic replacement. Intrinsic transmissibility also plays a role, with higher *R*_0_ promoting frequent emergence of antigenically novel strains [31, 30]. From a theoretical standpoint, successive strain replacement corresponds to epidemic dynamics that admit traveling wave solutions, in which infections create immunity that pushes existing strains in a single direction in phenotypic space [33, 24, 34, 35, 36]. These dynamics agree with representations of H3N2 antigenic evolution, with strains following a low-dimensional trajectory in a higher-dimensional antigenic space [13, 37].

A common assumption in these investigations is that individuals acquire immunity specific to any strain that infects them, regardless of infection history [20, 33, 22, 3, 23]. Decades of experimental and observational evidence of “original antigenic sin” (OAS) have demonstrated that secondary responses expand memory B cell responses that target previously encountered epitopes, limiting the generation of responses to new sites [38, 39, 40, 41, 42, 43, 44, 45, 46, 47, 48, 49]. This is especially apparent in responses to less immunogenic influenza vaccines [50, 51, 42]. Proposed mechanisms for this limited response to new sites include clearance of antigen by preexisting immunity [52, 53] and the reduced requirements for memory compared to naive B cell activation (e.g., [54, 55, 56]). A straightforward corollary is that no protection can be derived from nonexistent responses to new sites, and thus individuals infected or vaccinated with the same strain may be differently protected to other strains, depending on which epitopes they target [57].

OAS might sometimes boost antibody responses to a site that are cross-reactive, blunting new responses to that site, but are not necessarily so protective. Decoy non-neutralizing epitopes are a feature of some pathogens [58, 59], but in influenza, this discrepancy between reactivity and protection appears to arise occasionally from OAS. For instance, middle-aged adults infected with H3N2 sometimes boost antibody titers that bind well by ELISA but show negligible neutralization activity against the infecting strain [60]. Ferrets infected with one influenza A subtype boost their stalk antibodies to that subtype on later infection with another subtype, blunting new responses [61]. The same pattern appears in children infected with H1N1 before H3N2 [61]. These recalled antibodies appear to be very poor binders to the second subtype, and although the in vivo consequences of hundreds-fold reductions in affinity are uncertain [62], it is plausible they might be accompanied by reduced protection, as suggested by experimental infections in mice [63]. Thus, OAS might influence the protection derived from secondary exposures in two ways: first, by preventing any response to new sites, and second, by recalling memory responses to new but similar sites that poorly bind and weakly neutralize the new site. Either mechanism can in theory prevent the creation of protective memory to the infecting strain.

It is not known how OAS might affect pathogen evolution. Transmission models involving OAS have assumed that only one strain circulates at any time and always mutates between seasons [64, 65], and thus have not addressed how OAS impacts antigenic evolution. These transmission models predict that OAS impacts population immunity, leading to “immune blind spots” in different age groups [65], which might relate to recent evidence of ∼24-year cycles in the induction of strong antibody responses to H3N2 [66].

We investigate the consequences of OAS for pathogen evolution, and specifically for the conditions that permit successive strain replacement. Our mathematical models of host-pathogen coevolution incorporate OAS via the mechanisms described above. Both models allow for infection without creation of strain-specific memory through tunable parameters that allow cross-reactive responses to blunt the generation of new ones without conferring protection at that epitope. One model assumes a cross-reactive response at one epitope is sufficient to block the induction of new memory to the rest, and the other assumes that any epitope that is sufficiently diverged will elicit a new response. Nature is likely somewhere in between, depending on the type and number of secondary exposures [67, 68, 53, 63]. We show that these contrasting assumptions do not change our main result: OAS implies an upper bound on the basic reproduction number *R*_0_ of pathogens that can exhibit successive strain replacement, and this bound implies limits on the speed of evolution, the standing antigenic diversity, and the time to the most recent common ancestor in this regime. OAS thus narrows the conditions in which serial strain replacement is likely to occur.

## Model

We separately consider the dynamics of protection and memory creation in individuals before describing pathogen dynamics in the host population.

### Host-scale protection

Pathogen strains and strain-specific memories are parametrized as real vectors in an abstract, continuous *d*-dimensional antigenic space in which the distance between two points represents the extent to which a strain-specific memory at one point protects against infection with a strain at the other point, as detailed mathematically below. The model assumes that each of the *d* dimensions of antigenic space corresponds to a physical region bound by antibodies, and the footprints of antibodies binding this region are entirely contained within it. We refer to these antibody-binding regions as epitopes. A similar model with each axis corresponding to one epitope was used in [69].

Individuals infected with a strain of the pathogen are also infectious. An infected individual can transmit either the infecting strain or a mutant strain. Infected individuals expose on average *R*_0_ other individuals during their infection, where *R*_0_ is the basic reproduction number and does not vary by strain. The duration of infection also does not vary by strain.

Individuals are born without immunological memory. A naive individual always becomes infected when exposed to a strain and forms strain-specific memory on infection. A non-naive individual who becomes infected might develop memory specific to that strain. The set *𝒮* of strains 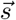 to which an individual has strain-specific memory at time *t* is their memory profile at *t*. An individual has at most one specific memory targeting (i.e., derived from infection with) each strain. The number of strains in *𝒮* equals *m*. In general, the number of strains in memory, *m*, would be expected to vary among individuals, though the population-scale model will assume that a fixed value of *m* is a reasonable approximation for computing protection.

Memory decreases the probability of infection on exposure. Individuals cannot be reinfected by strains in their memory profile. Memory is cross-protective, meaning that strain-specific memory decreases the probability of infection with similar strains (Fig. 1). Cross-protection declines exponentially with the Euclidean antigenic distance between the strain in memory 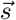 and the challenge strain 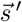,

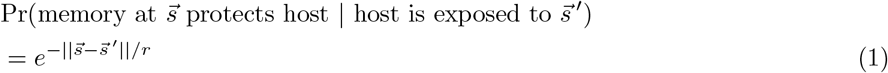

with 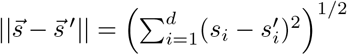. The quantity *r* is the cross-protection distance. It is an independent parameter that determines how antigenic change affects the protectiveness of existing memory to related strains. If *r* is high, cross-protection is high, and large mutations in antigenic space are required for protection to drop.

**Figure 1.**
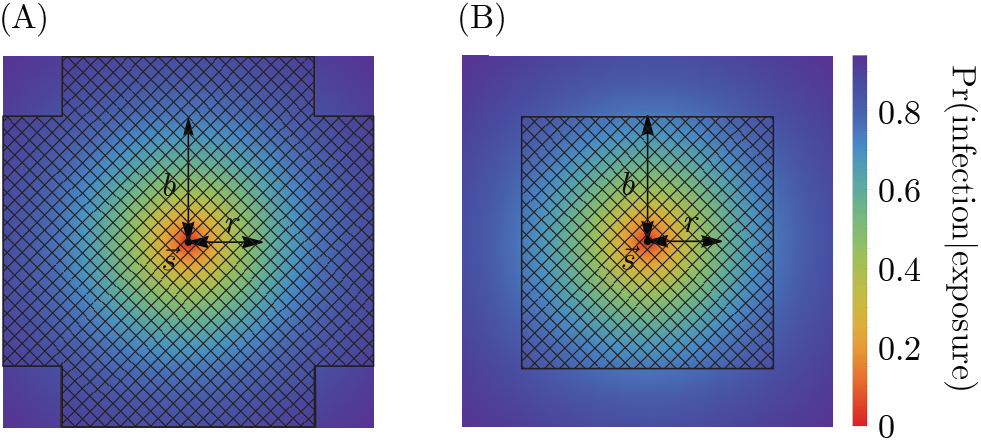
Cross-protection and blunting from a single memory in 2-dimensional antigenic space. A strain-specific memory 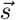 at the center protects against infection by nearby strains according to Eq. 1. The distance over which protection drops by a factor of 1/e is *r*; the blunting distance is *b*; hatched regions represent excluded regions in the (A) every- and (B) any-epitope models. Only infections with strains outside these regions would generate new memory. Here, *b* = 1.7 and *r* = 1.25.

Eq. 1 defines the probability that a single strain-specific memory protects against infection by another strain. Each strain 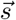 in an individual’s memory profile *𝒮* contributes to protection against a challenge strain 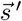. The probability of infection for an individual with memory profile *𝒮* exposed to a challenge strain 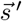 is given by the product of the probabilities of each memory separately failing to protect,

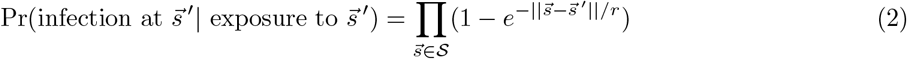

Implicit in Eq. 2 is the assumption that protection against infection depends on the unordered set of strains in the memory profile, yet we have claimed that OAS induces a dependence on the order of exposure. Order dependence arises from the dynamics of memory creation.

### Creation of new memory

Infections and vaccinations do not always generate new specific memory to the challenge strain. Whether an individual generates a strain-specific memory depends on their existing memory profile. We model OAS via blunting, and consider two models, the “every-epitope” model and the “any-epitope” model, corresponding to the limits of biologically plausible scenarios. In each, a blunting distance, *b*, sets a threshold antigenic distance for developing new memory.

In the every-epitope model, the challenge strain 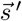 must escape immunity at every epitope to create strain-specific memory (Fig. 1); escape at only some epitopes can be overcome by adaptive immune responses, such as antibodies, binding conserved regions [69]. More precisely, an individual challenged with a strain 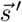 creates strain-specific memory if 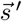 differs from every strain 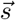 in their memory profile *𝒮* by an antigenic distance of at least *b* in each of its *d* antigenic space coordinates,

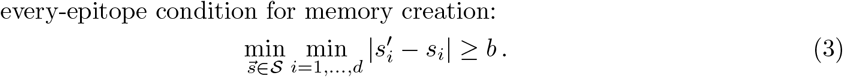

In the any-epitope model, it is enough for the challenge strain 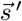 to escape immunity in a single epitope to create strain-specific memory (Fig. 1) [3]. An individual challenged with a strain 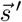 creates strain-specific memory if 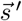 differs from every strain 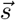 in their memory profile *𝒮* by at least *b* in at least one epitope (which may not be the same epitope across *𝒮*),

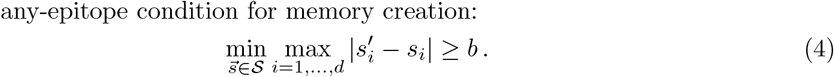

These models can be conceptualized as different types of exposures. The every-epitope model could represent an exposure to a small dose of antigen, for example via natural infection, where the presence of any neutralizing memory can limit pathogen replication, reducing antigen availability enough to prevent the generation of new memory. The any-epitope model could represent an exposure to a high dose of antigen or an immunogenic vaccine that forces a response to diverged epitopes even in the presence of memory that would ordinarily suppress a new response. This effect has been shown in animal models since the earliest observations of OAS [46], and more recently in molecular detail in mice [68, 48].

Throughout the remaining analysis, we assume that immune memory can cross-react with a larger set of strains than it can protect against (*b* > *r*). That is, it can suppress the formation of new memory against some infecting strains. While it is not biologically impossible that *r* would be greater than *b*, this would require enough antigen to provoke a memory response without causing an infection. This might occur through vaccination but will not be explored here.

### Population-scale infection-immunity coupling

At the population scale, our model is equivalent to the model of pathogen-immune coevolution developed in [36] with the added assumptions that each dimension corresponds to an epitope in the proper coordinates and that each epitope antigenically evolves at a rate that does not depend on time. The population-scale description is formulated in terms of two densities in antigenic space: the total number of individuals infected by each strain 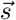 at time *t* and the total number of individuals with strain-specific memory to 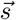 at time *t*, denoted 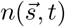 and 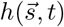, respectively. In transitioning to a density-based approach, we trade complete information about each individual’s memory profile for tractability. This is similar to the choice of a previous cohort-level description for studying OAS [65], although the motivation is different as we do not incorporate age structure.

Strain fitness is determined by the number of new infections per infectiousness period, which depends on individuals’ immunity. The expected protection against infection upon exposure to strain 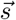 conferred by a single memory chosen uniformly at random from 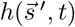 is denoted 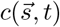. Using the definition in Eq. 1, this is equal to

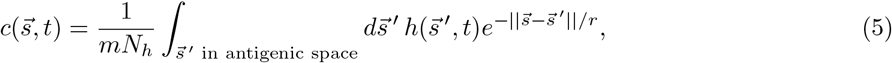

where *N*_*h*_ is the size of the population and the factor *mN*_*h*_ adjusts for the normalization of 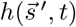. In analogy with Eq. 2, we have

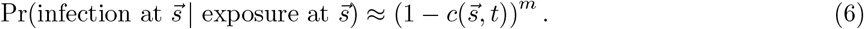

Equations 5 and 6 rely on an assumption that the average protection against infection across individuals is well approximated by the average protection in a system where the 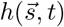 memories are distributed randomly among individuals, with each individual having the same number of memories, *m* (see Methods for further discussion on this point). Defining *R*_0_, the basic reproduction number of the pathogen, to be the expected number of people an infected individual encounters during their infection and would infect (assuming complete susceptibility), the fitness of strain 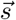 at time *t* is

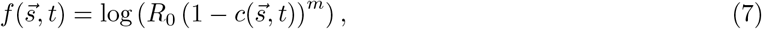

as in [36].

We model the successive strain replacement regime using the traveling wave model described in [36]. Traveling wave solutions arise from stochastic coupled dynamical equations between the infection density and the memory density that take into account fitness differences among strains, mutations, noise in the transmission process, and memory addition and deletion:

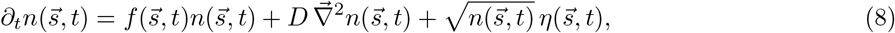

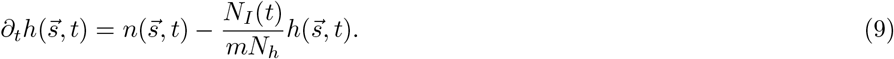

The first equation (Eq. 8) states that as the pathogen spreads, each strain grows or shrinks according to its fitness (Eq. 7); mutations are frequent and have small effect, such that the diffusion approximation is appropriate with diffusion constant *D*; and pathogen spread is stochastic, with Gaussian process noise, where 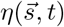 is a unit Gaussian 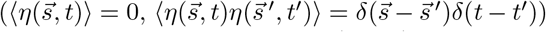, and the prefactor 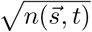 sets the mean and standard deviation. The second equation (Eq. 9) states that the change in memory at a given time is determined by memory addition at the locations of infections and by memory deletion uniformly at random: each person acquires a new strain-specific memory at 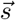 when they are infected by strain 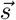, and at that time, they lose one memory from their memory profile *𝒮*, such that each person maintains a fixed number of *m* memories over time. Time is modeled in units of the duration of infection.

### Successive strain replacement dynamics

Successive strain replacement dynamics are conceptualized as an infection distribution that takes the form of a localized clump that maintains its shape as it moves through antigenic space at a constant speed and direction and is Gaussian in the direction of motion to reflect the assumption that the mutation process is diffusive. The localization implies that strains are immunologically well-defined, with low antigenic diversity around any point at any time. The constant speed and direction amount to assuming that the effect of stochasticity in orthogonal directions is negligible. While the speed at which the wave travels is a derived quantity determined by the transmission and immunity parameters governing the dynamics, its direction of travel is determined by the relative rates at which different epitopes antigenically evolve, an independent input to the system which we parameterize by the angles between the direction the wave is traveling and the axes for individual epitopes, *θ*_1_, …, *θ*_*d*_ (Fig. 2C).

**Figure 2.**
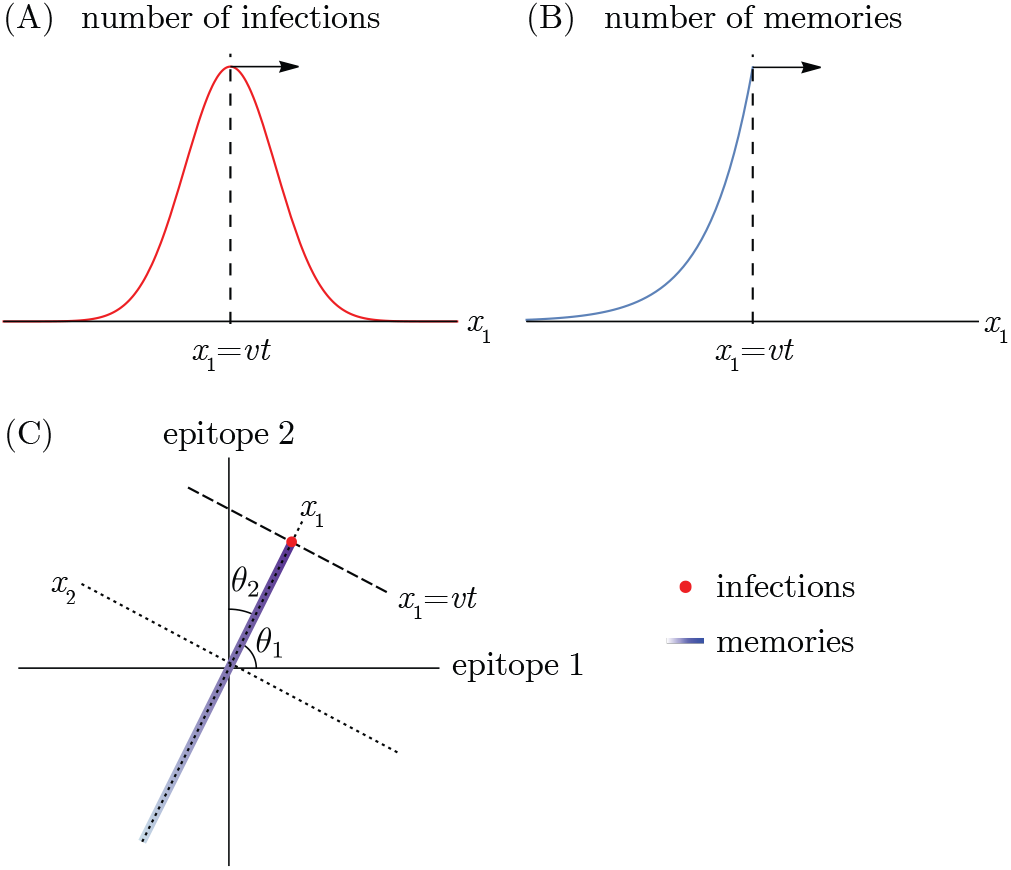
Successive strain replacement as traveling waves in antigenic space. (A) Strain replacement is modeled as a localized infection distribution moving along a one-dimensional trajectory in *d*-dimensional antigenic space with constant velocity 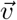. (B) The memory distribution is an exponentially decaying distribution with wavefront at *x*_1_ = *vt*, localized in orthogonal directions. (C) The axes aligned with the direction of motion of the infection and memory distributions are not the same as the epitope-aligned axes. The localized infection peak is visualized as a point at the location of the wavefront.

Successive strain dynamics are most simply expressed in coordinates where one axis is aligned with the direction of motion, with the remaining axes aligned orthogonally. We use 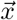 (rather than 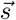) to indicate “wave-adapted” antigenic space coordinates where the *x*_1_ axis lies along the axis of antigenic evolution.

As in [36], the infection distribution takes the form

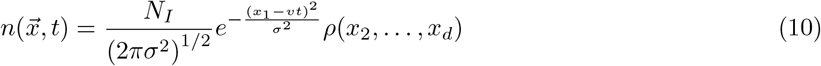

where *N*_*I*_ is the number of infected individuals, *v* is the wave velocity, *σ* is the standard deviation of the Gaussian in the direction of motion, and *ρ* is a normalized distribution. Both the Gaussian and *ρ* are taken to be localized to a region of linear dimension much less than the cross-protection distance *r* (*σ* ≪ *r*) (Fig. 2A). We are interested in the case where the cross-protection distance is less than the blunting distance, and thus these are also localized to a region of linear dimension much less than the blunting distance *b* (*σ* ≪ *b*).

When the pathogen undergoes serial strain replacement, represented by the infection distribution in Eq. 10, the population develops a memory distribution in the shape of a wavefront with maximum at the peak of the infection distribution that falls off exponentially fast in the direction of earlier infections (Fig. 2B). There is no specific memory ahead of the peak of infections. The wavefront travels at the same speed, *v*, as the infection distribution, and in the same direction. Since in this model each person has *m* memories in their memory profile, the total number of memories in the population (the integral over the entire memory distribution) equals the size of the population, *N*_*h*_, times *m*.

As derived in [36], the speed of antigenic evolution, *v*; standard deviation of the infection distribution, *σ*; and number of infected individuals, *N*_*I*_, are not independently specified. Given the dynamics specified by Eqs. 8 and 9, they are determined by the cross-protection distance, *r*; the number of individuals in the population, *N*_*h*_; the diffusion coefficient characterizing the effect of mutations, *D*; the number of memories per person, *m*; and the basic reproduction number, *R*_0_. See Methods for the relevant expressions.

## Results

We first describe static properties that the population memory and protection distributions must satisfy to be consistent with the two blunting models, regardless of the transmission dynamics that give rise to them. We then derive the consequences for pathogen evolution: to obtain serial strain replacement, the speed of evolution, *v*; the strain diversity, *σ*; and the time to most recent common ancestor, *t*_MRCA_, are constrained by an upper bound on the basic reproduction number, *R*_0_. This upper bound is determined by the characteristic distances defining memory creation and cross-protection (*b* and *r*, respectively), and by how changes in individual epitopes impact protection (the blunting model).

### Blunting neighborhoods

In both the every-epitope model and the any-epitope model, blunting implies a geometric condition that no individual’s memory profile contains specific memory to strains that are too close to one another. The precise definition of “too close” differs between the models.

The every-epitope assumption (Eq. 3) implies that for every pair of strains 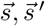 in a host’s memory profile *𝒮*, the difference 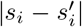 is at least *b* in each dimension *i* = 1, …, *d*. This means that given any reference point 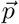 in antigenic space, for any dimension *i*, there exists at most one strain in *𝒮* that is a distance less than or equal to 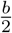 from 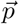 in that dimension. The set of all such strains defines *d* stripe-shaped regions in which there can be at most one memory: the set of points 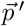 with 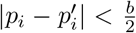 for at least one value of *I* (Fig. 3A). Each stripe has width *b* parallel to the *i*th axis and extends infinitely in the remaining (*d* − 1) axes (derivation in Methods). We refer to these stripes as the blunting neighborhoods of 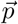 in the every-epitope model and denote them 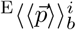.

**Figure 3.**
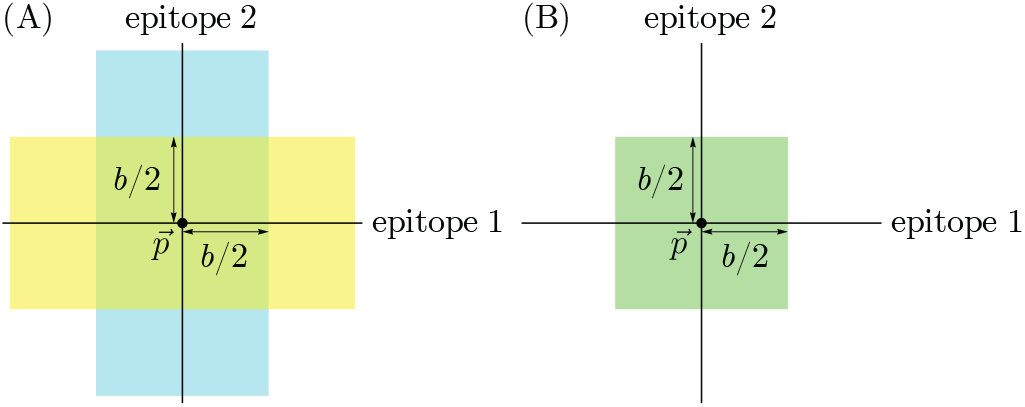
Blunting neighborhoods about reference point 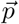 for two blunting models, shown in 2*d* antigenic space. If an individual has memory to a strain in a blunting neighborhood of a given reference point and is later infected by a different strain in that neighborhood, no new memory will be created. (A) Blunting neighborhoods for epitope 1 (yellow) and epitope 2 (blue) about 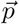 in the every-epitope model. (B) Blunting neighborhood about 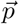 in the any-epitope model.

The any-epitope assumption (Eq. 4) implies that for every pair of strains 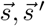 in a host’s memory profile *𝒮*, the difference 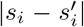 is at least *b* in at least one dimension *i* = 1, …, *d*. This means that given any reference point 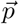 in antigenic space, there exists at most one strain in *𝒮* that is within the distance 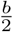 from 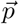 in all dimensions, defining a *d*-dimensional cube with side length *b* centered at 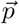 (Fig. 3B) (derivation in Methods). We refer to this cube as the blunting neighborhood of 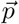 in the any-epitope model and denote it 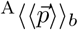.

### Order-dependent protection

Both proposed blunting models are consistent with a major observation motivating our study of OAS: they imply that individuals’ immunity depends not only on the set of strains to which they have been exposed but also on the order of exposure (Fig. 4) [70, 60].

**Figure 4.**
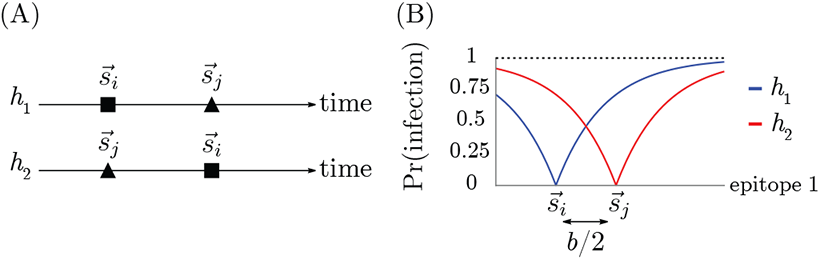
Protection is order-dependent with blunting. (A) Infection history of hosts *h*_1_ and *h*_2_. (B) Memory profiles and protection distributions of hosts *h*_1_ and *h*_2_. Here, 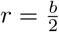.

Consider the simplest case of a pathogen whose evolution takes place in one-dimensional antigenic space. In one dimension, the every-epitope model and the any-epitope model are equivalent. Two strains are present: one, 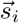, at the origin and the other, 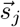, at *b*/2. Consider two different hosts, *h*_1_ and *h*_2_, both of whom have been infected by strains 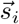 and 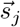 but in different orders: *h*_1_ was infected by 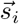 followed by 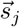, and *h*_2_ was infected by 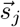 followed by 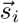 (Fig. 4A). Host *h*_1_ develops specific memory to strain 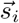 but not to strain 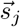, whereas host *h*_2_ develops specific memory to strain 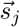 but not 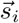 (Fig. 4B). Given these memory profiles, the probability that each host is infected by any strain 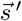 is given by

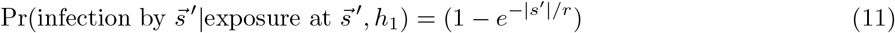

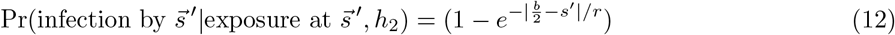

For example, if host *h*_1_ is exposed to the strain 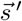 at *b*, they will be infected with probability (1 − *e*^*−b/r*^), whereas if host *h*_2_ is exposed to the same 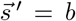, they will be infected with probability (1 − *e*^*−b/*2*r*^), a smaller value. Although *h*_1_ and *h*_2_ have the same set of exposures in their history, blunting dynamics lead to *h*_1_ having less protection to 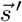 than *h*_2_ (Fig. 4B). Both blunting models therefore predict order-dependent protection to antigenically diverged strains.

In higher dimensions, the mechanism by which the every- and the any-epitope model exhibit order-dependent protection is essentially the same as in one dimension, but because blunting neighborhoods are infinitely extended in the every-epitope model and localized in the any-epitope model, they differ in their implications for heterogeneity of protection in the population. In the every-epitope model, order dependence can induce strong heterogeneity of protection, as a pathogen with some slowly-evolving (i.e., conserved) epitopes and at least one fast-evolving epitope travels far before exiting blunting neighborhoods in the conserved direction. Since individuals do not create new memory until the pathogen exits the blunting region, and protection decreases exponentially with distance between the memory and challenge strains, experienced individuals will be less protected to circulating strains than individuals with more recent primary infections. By contrast, in the any-epitope model, order-dependence can have less impact over time, as there is a maximum distance the pathogen travels before leaving blunting neighborhoods. This means that two individuals’ protection cannot diverge arbitrarily as the pathogen evolves, since new infections will trigger memory creation more frequently even in experienced individuals. This distance is maximized when all epitopes evolve at similar rates.

### Immune blind spots

Epidemiological observations indicate that individuals do not always develop specific memory against a strain 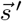 even after repeated exposures [71, 60]. Those strains occur in “immune blind spots” [65], which we show here create blind spots in protection as well. In both the every-epitope model and the any-epitope model, immune blind spots arise when *b* is large compared to *r*: the blunting conditions guarantee the existence of points that are too antigenically similar to existing memory to trigger creation of new memory upon exposure but are nonetheless far enough to escape cross-protection from infection. Intuitively, this is because OAS prevents memory profiles from becoming too dense, and thus limits the antigenic “coverage” of protection.

To demonstrate the existence of immune blind spots, we will consider a scenario for each model. In the every-epitope model, any strain 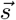 in the memory profile defines an infinitely extended *d*-dimensional cross-shaped region where no additional memory can be created (Fig. 5A). For example, any strain 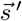 sharing at least one epitope with 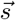 can never be added to the memory profile. Suppose strain 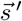 has all but one epitope conserved with 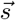, and the remaining epitope is so far diverged as to escape all existing memory (i.e., the distance in that direction is much greater than both the cross-protection distance and the blunting distance). Since cross-protection to 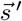 depends on the Euclidean distance from 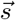, the diverged epitope prevents 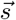 from protecting against 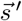 despite shared epitopes. In fact, if the blunting distance is much greater than the cross-protection distance, an individual with 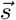 in their memory profile will *never* develop protection to 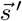. This is because future exposures will create memory only to strains diverged from 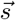 by at least the blunting distance in every direction, and if *b* ≫ *r*, such strains cannot protect against strains sharing epitopes with 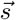, including 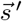 (Fig. 5A). When the blunting and cross-protection distances are comparable, a strain diverged from 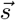 (and therefore 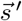) by exactly the blunting distance in the conserved directions can partially cross-protect against 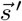. This means that the immune blind spot disappears if the blunting distance *b* is sufficiently close to the cross-protection distance *r*. An explicit calculation of the loss of protection at 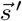 as a function of the ratio of blunting to cross-protection (Methods) shows that the immune blind spot appears at modest values of *b*/*r*, and the approximate minimum threshold for the appearance of immune blind spots shrinks as the number of epitopes (*d*) increases (Fig. 5C).

**Figure 5.**
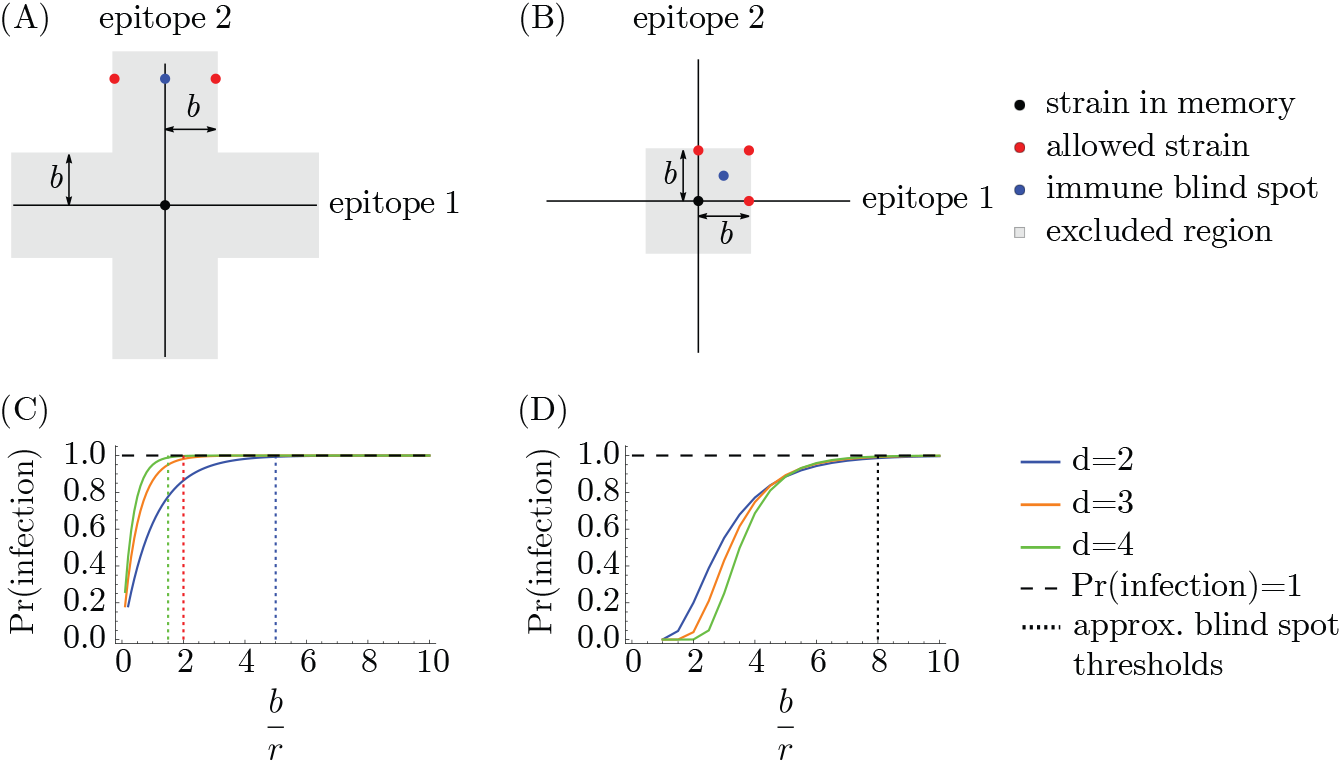
Immune blind spots arise when *b* is large compared to *r*. Given a memory at the origin, the “allowed strains” are examples of the closest possible strains that could be added to the memory profile. The protection is necessarily low at the immunity blind spot strains regardless of infection history in (A) the every-epitope model and (B) the any-epitope model as long as 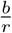 is sufficiently large (and, in (A), the distance from the memory strain to the immune blind spot is greater than both the blunting distance and the cross-protection distance). (C, D) show the probability of infection at the blind spot points considered here for (C) the every-epitope model and (D) the any-epitope model. Vertical dotted lines indicate approximate values of *b*/*r* above which infection is essentially guaranteed (decreasing with dimension in (C), dimension-independent in (D)).

Similarly, in the any-epitope model, any strain 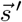 that is diverged by less than the blunting length From 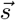 in at least one epitope cannot be added to the memory profile, and individuals will maintain some susceptibility to that strain as long as *b* > *r*. One example is a strain 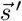 which is diverged from 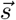 by antigenic distance of half the blunting length (*b*/2) in every epitope (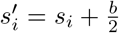, for example) (Fig. 5B). The closest strains to 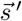 that could be added to the memory profile are strains that are diverged from the reference strain 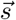 in at least one epitope by exactly the blunting distance (+*b*). The strain 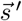 escapes cross-protection if 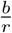 is very large, but may be protected from these nearby strains if the cross-protection distance is comparable to the blunting distance (Fig. 5D). As long as 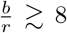, infection upon exposure to strain 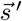 is essentially guaranteed, and so 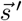 falls in an immune blind spot (Fig. 5D) (Methods).

### Population-scale memory constraint

Both blunting models imply maximum total numbers of specific memories the population maintains in certain regions of antigenic space. This follows from the existence of blunting neighborhoods. Each host has at most one memory within the blunting neighborhood(s) 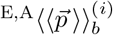 of any reference point 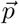 in antigenic space. (The *i* index applies to the every-epitope model only.) It follows by taking the sum of memories at each point over all hosts that in any collection of *N*_*h*_ non-naive individuals with collective memory distribution 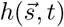, the total number of memories in each blunting neighborhood of 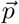 cannot exceed *N*_*h*_. In symbols, for both blunting models, we have

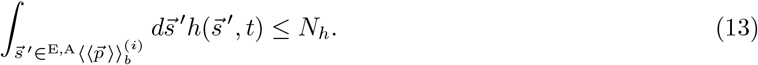

The every-epitope model and the any-epitope model differ in the regions over which the integral is taken, with the every-epitope model integrating over *d* infinitely extended stripes to 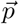, and the any-epitope model integrating over a single *d*-cube to 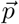 (Fig. 3).

We refer to Eq. 13 as the OAS constraint. It holds in any model of OAS with blunting.

### Limits on successive strain replacement dynamics

We find that the conditions for serial strain replacement are harder to obtain in models that assume blunting (OAS). Only pathogens with a sufficiently low *R*_0_ compared to the ratio of the cross-reactivity distance to the blunting distance (formally, *e*^*b/r*^) can exhibit successive strain replacement dynamics in populations where individuals accumulate many immune memories (i.e., in the large-*m* limit).^1^ This bound implies a maximum speed of antigenic evolution that depends on the blunting model.

In the dynamics described by [36], the population accumulates memories in a small interval close to the wavefront. This localized buildup of memory is necessary for maintaining the fitness gradient required to force the infection distribution to move at a constant speed while maintaining its shape. We have shown, however, that the OAS constraint prevents arbitrarily localized buildup of memory. Since the shape of blunting neighborhoods depends on the blunting model, the amount of memory close to the wavefront is also sensitive to specific assumptions about blunting. Specifically, OAS prevents memory accumulation in an interval whose length is determined by the projection of the direction the wave travels in the fastest (every-epitope model) or slowest (any-epitope model) direction (Methods). We refer to the angle between the fastest (slowest) direction in the every- (any-) epitope model as *θ*^*^.

The OAS constraint requires that

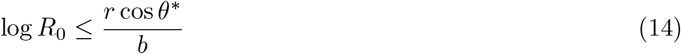

in the large-*m* limit (Fig. 6; derivation in Methods). Since cos *θ*_*i*_ measures how quickly the *i*th epitope evolves compared to the other (*d* − 1) epitopes, this equation says that successive strain replacement is possible only if *R*_0_ is not too large compared to the inverse of the blunting distance *b* in units of the cross-protection distance *r*, modulated by the projection onto the most important epitope for that model.

**Figure 6.**
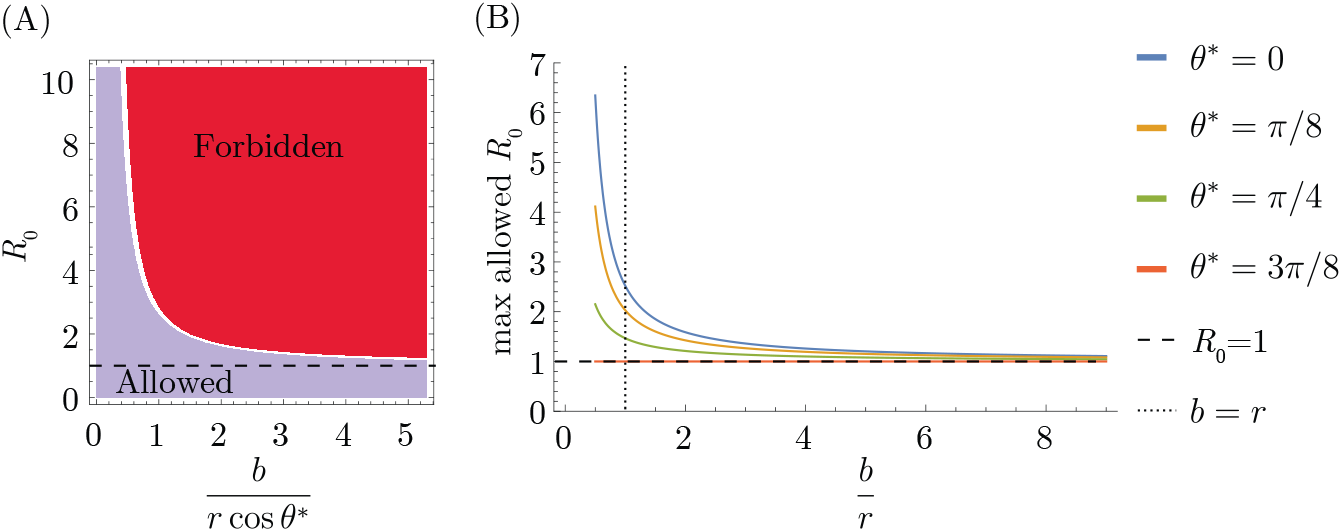
Transmission limit implied by the OAS constraint. (A) Strain replacement imposes a maximum value on *R*_0_. (B) Maximum value for *R*_0_ as a function of 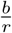 for different values of *θ*^*^.

In the traveling wave model, the cross-protection distance, *r*; the population size, *N*_*h*_; the diffusion constant, *D*; and the basic reproduction number, *R*_0_ govern pathogen evolution, including the speed *v* at which the pathogen evolves, the standard deviation *σ* of infections in the direction of evolution, and the time to most recent common ancestor, *t*_MRCA_, of cocirculating strains [36]. The inequality in Eq. 14 implies that blunting leads to (*N*_*h*_- and *D*-dependent) maximum *v*, minimum *σ*, and minimum *t*_MRCA_ in the context of successive strain replacement (Fig. 7). These limiting values are achieved for pathogens that saturate the inequality (Methods). For example, if the blunting length *b* is equal to twice the cross-protection length *r* and there are two epitopes evolving at the same rate (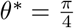 for both blunting models), then *r* cos *θ*^*^/*b* ∼ 0.35, and the maximum *R*_0_ consistent with successive strain replacement dynamics is ∼ 1.42. In this case, the minimum time required for the viral population (peak of the infection distribution) to travel the cross-reactivity distance varies up to ∼ 100 times the duration of infection (for, e.g., *r* = 5); the minimum standard deviation of the viral population varies between ∼ 0.1 and 0.5 cross-protection distances for values of *r* between 1 and 5; and the minimum time to the most recent common ancestor for two cocirculating strains varies between ∼ 100 − 400 times the duration of infection for values of *r* between 1 and 5.

**Figure 7.**
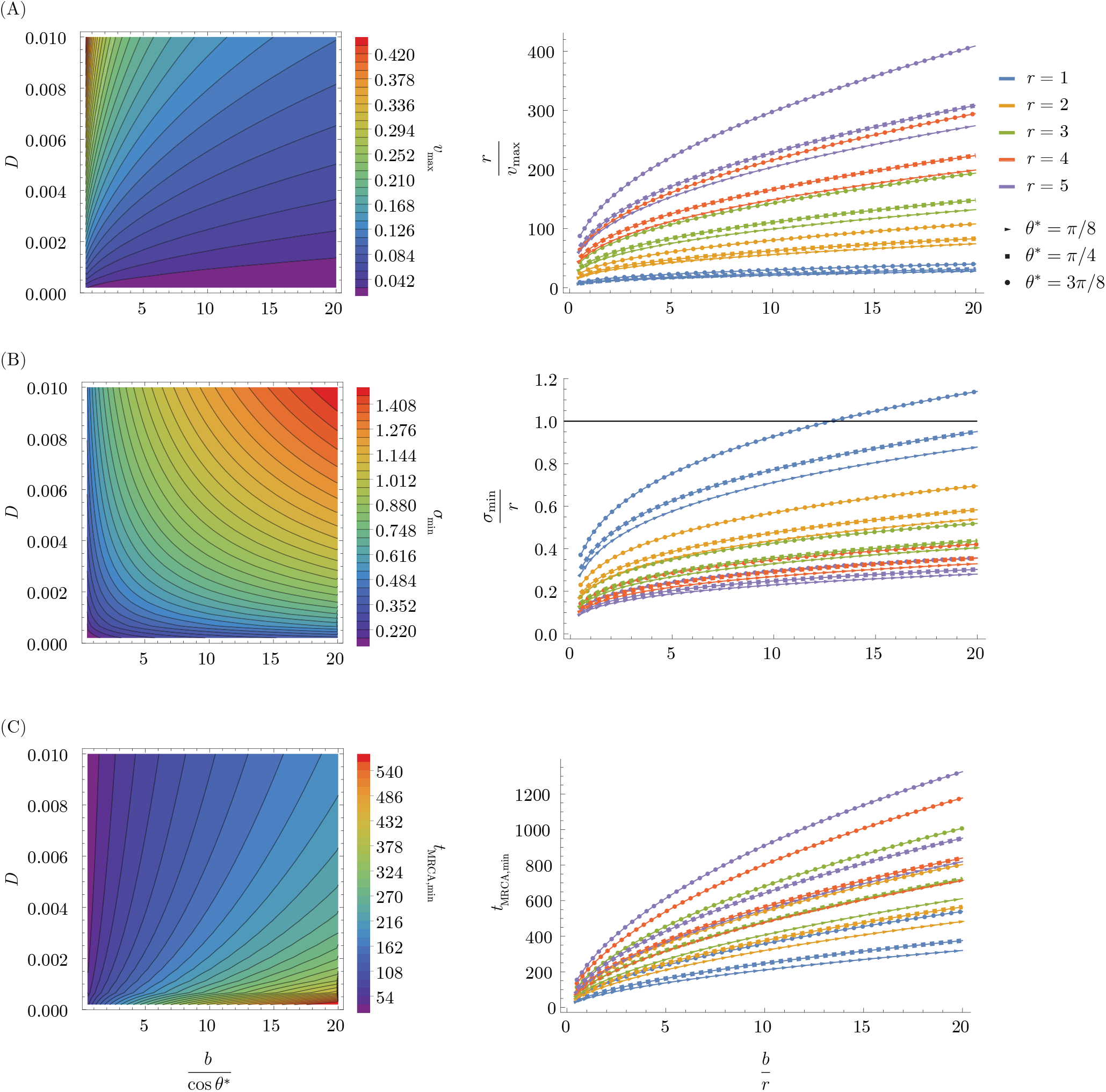
Dynamical limits implied by the OAS constraint. (A) Maximum speed of evolution *v* (antigenic distance per infection), which depends on the diffusion constant *D* and the blunting distance modulated by the projection onto the fastest (slowest) evolving epitope for the every (any) epitope model (left), and the time for the infection peak to travel the cross-protection distance as a function of 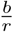 for different values of *r* and *θ*^*^ at saturating value of *v*_max_(right). (B) Minimum standing diversity *σ* (antigenic distance) as a function of the diffusion coefficient and blunting distance modulated by the projection onto the fastest (slowest) evolving epitope for the every (any) epitope model (left), and the minimum standing diversity in units of cross-protection distance as a function of 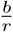 for different values of *r* and *θ*^*^. Parameter combinations lying above or close to the black solid line at *σ*_min_/*r* = 1 cannot lead to serial strain replacement, as the traveling wave solution requires *σ*/*r* ≪ 1. (C) Minimum time to most recent common ancestor, *t*_MRCA_ (measured in duration of infection), as a function of the diffusion coefficient *D* and the blunting distance modulated by the projection onto the fastest (slowest) evolving epitope for the every (any) epitope model (left), shown as a function of *b*/*r* for different values of *r* and *θ*^*^ (right). Here, *N*_*h*_ = 10^9^ (left and right panels) and *D* = 0.004 (right panels).

## Discussion

Many factors have been hypothesized to lead to serial strain replacement, but theoretical tests of these hypotheses have tended to assume that individuals uniformly develop protective strain-specific immunity from infection. We have shown that OAS limits the density of population memory in antigenic space, a picture we expect to hold broadly across models of strain evolution. This bound on the number of memories constrains the successive strain replacement regime to low-transmissibility pathogens, with “more OAS”—a greater ability of memory to blunt the formation of new responses—corresponding to a tighter bound. Our results imply that observed patterns of serial replacement in nature may be less stable than thought.

This work represents a qualitatively new approach compared to past studies of successive strain replacement. Previous work on waves in antigenic space [36] found that traveling waves solve the model for any set of parameters. Other investigations into the existence of traveling waves in pathogen evolution [24, 72, 73, 74] focused on phenomena such as disease treatment [73, 74] without including OAS. These models found traveling wave solutions for all *R*_0_ > 1 with a *minimum* speed of evolution, as opposed to our result, which implies that OAS imposes a *maximum* speed of evolution. The previous studies that implemented OAS assumed successive strain replacement and examined its impact in age-structured models [64, 65]. Our results are complementary, in that we identify where successive strain replacement is likely to arise in the first place.

By preventing arbitrary buildup of population memory in localized regions of antigenic space, OAS effectively limits the steepness of the fitness landscape that drives selection to escape immunity, and thus restricts the space of possible infection dynamics. Traveling waves are one example of a dynamical pattern that is subject to this restriction, as they require a dense buildup of memory near the wavefront. The necessary density depends on the characteristic cross-protection distance *r* and the basic reproduction number *R*_0_, with smaller *r* and larger *R*_0_ resulting in more memory near the wavefront. Since the blunting distance *b* controls the number of memories that are “captured” in the blunting neighborhood at the wavefront, traveling wave dynamics are only consistent for small enough *b* in relation to *r*^*−*1^ and *R*_0_. If the OAS bound is not satisfied, then the wave solution is unstable to the frequent generation of escape mutants. Depending on the diffusion coefficient, this could lead to the wave spreading or to continual generation and disappearance of multiple peaks, similar to the mound and comb patterns described in a similar model of virus-immune coevolution [75].

The implementation of the OAS bound in terms of blunting neighborhoods also demonstrates an intimate connection between OAS and the rules of memory creation in the presence of multiple evolving epitopes. The geometry of blunting neighborhoods runs parallel to the epitope-aligned axes. By contrast, as long as more than one epitope is evolving, the traveling wave does not travel parallel to any axis. Consequently, the number of memories captured in the blunting neighborhood near the wavefront depends on the relative speeds at which different epitopes evolve. Because the shape of blunting neighborhoods depends on the interactions among epitopes, the slowest evolving epitope determines the upper bound on *R*_0_ for the every-epitope model, and the fastest evolving epitope determines the bound for the any-epitope model.

The importance of *R*_0_ and epitopes’ rates of evolution suggests that effective public health measures, including non-pharmaceutical interventions and vaccines, might make pathogens more likely to exhibit successive strain replacement. An immunogenic, well-matched vaccine can deliver a high enough or “long enough” dose of antigen to overcome immune memory, forcing responses to new sites and the creation of strain-specific memory (e.g., [68, 48, 53]). In contrast, at small doses of antigen, immune memory might effectively clear antigen unless every epitope is so antigenically diverged that new strain-specific memory can form. This suggests that population immunity dominated by immunogenic vaccines is likely better approximated by the any-epitope model. Since cos *θ*^*^ is always greater in the any-epitope model than in the every-epitope model, the OAS bound on *R*_0_ is weaker. In other words, we expect pathogens will be more likely to exhibit successive strain replacement and less likely to diversify when vaccines can induce broader memory than infections.

One limitation of this work is that the population-level OAS constraint is imposed, rather than allowed to emerge from the individual-level model dynamics. Although we expect successive strain replacement models to generically induce memory buildup near the wavefront, we would need to incorporate additional mechanisms into the model for memory dynamics to recapitulate OAS endogenously. We still expect that regimes with a moving wavefront exist, but their functional form may be more complex or even time-dependent. Since all solutions to such a model would satisfy the OAS constraint by design, restrictions on the parameter regimes where successive strain replacement occurs would derive from existence conditions on wavelike solutions to the differential equation rather than consistency conditions on memory at the wavefront. Since in such a model memory could saturate, we hypothesize that a cross-protective, short-term immunity might be required to avoid repeated infections from accumulating, consistent with the conditions in [12, 27]. With dynamics that enforce OAS, evolutionary dynamics could be explored in more regimes, e.g., to identify the conditions under which branching occurs. It would be interesting to compare such a study to [34, 6], which construct similar phase diagrams in models without OAS.

There is substantial heterogeneity in antibody-mediated protection against influenza and SARS-CoV-2 in the human population, with individuals showing diversity in their antibody landscapes [76, 39, 66, 77] and the particular epitopes they target, including their sensitivity to different escape mutations [78, 79, 80, 81, 71]. Some of this heterogeneity arises from differences in initial exposures [71], but there is striking diversity within birth cohorts [77, 66]. Future versions of this model could include different subpopulations, distinguished by their memory repertoires, that are tracked over time and parameterized empirically.

Several assumptions should be reexamined in applying this theory to data, namely, the constant cross-protection distance *r* across epitopes, the simple memory creation and cross-protection functions, the single dimension of antigenic space for each epitope, and the artificial dichotomy of the any-epitope and every- epitope models. The correlates of protection for pathogens including influenza and SARS-CoV-2, necessary for understanding the cross-protection distance *r*, are not well understood and require additional investigation via prospective studies and challenge experiments. Other model assumptions, especially the blunting distance *b* and any- and every-epitope model forms, require a more quantitative understanding of the dynamics of immune memory. Longitudinal studies that combine diverse exposure histories with detailed immune profiling will help to accelerate understanding on these fronts.

## Methods and derivations

### More details on serial strain replacement assumptions

We expand on the assumptions underlying the serial strain replacement model, which are nontrivial for a model of OAS and trade full information on the individual scale for tractability on the population scale. Given complete information about the memory profile of every individual, an individual’s protection against a strain 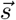 could be calculated, but with only the density 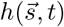 available, we must approximate the distribution of protection. We make a homogeneity assumption, i.e., that the average protection against infection is well approximated by the average protection in a system where the 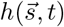 memories are distributed randomly among individuals, with each individual having memory to a fixed number of strains, *m*.

The homogeneity assumption precludes direct consideration of dynamics where the population consists of groups of individuals with highly correlated infection histories that differ substantially between groups, such as birth cohorts; however, this kind of structure could be accommodated by introducing a separate memory density for each cohort and specifying transmission dynamics between cohorts.

Similarly, we assume that the average susceptibility is well approximated by the susceptibility when each individual’s memory profile is an independent draw of *m* memories from 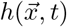, meaning that homogeneity extends to the number of exposures to the pathogen over time (and therefore over antigenic distance).

When the infection distribution 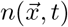 takes the localized form in Eq. 10 and the memory and infection distributions satisfy the coupled differential equations Eqs. 8 and 9, it is possible to write down an explicit equation for the form of 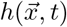. In a zoomed-out limit where *x*_1_ ≫ *σ* and *x*_*i*_ ≫ *b* for *i* = 2, …, *d*,

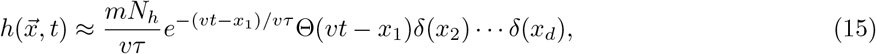

where *N*_*h*_ is the population size, *τ* = *mN*_*h*_/*N*_*I*_, and Θ is a step function, Θ(*y*) = 1 for *y* ≥ 0 and Θ(*y*) = 0 for *y* < 0. Eq. 15 follows from Eqs. 8 and 9, assuming 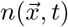 takes the form in Eq. 10 [36].

### Derivation of blunting neighborhood conditions

We derive the existence of blunting neighborhoods (Fig. 3) from the conditions for the creation of memory in the every-epitope and any-epitope models (Eqs. 3 and 4).

First, the every-epitope model: Let 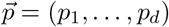 be a reference point in antigenic space. Suppose the strain 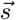 is in *𝒮*, and there exists at least one *j* for which 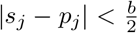. Suppose the host is infected by (or vaccinated with) strain 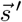 that is also within a distance 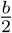 from 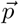 in dimension *j*. If both *d*_*j*_ ≡ (*s*_*j*_ − *p*_*j*_) and 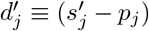 have the same sign, then 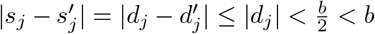, and therefore 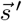 is not added to *𝒮*. If (*s*_*j*_ − *p*_*j*_) and 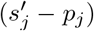 have different signs, then 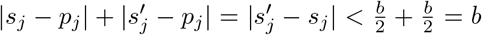, and so 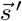 is not added to *𝒮*. Either way, 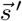 will not be added to *𝒮*, meaning that 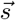 will remain the only strain in *𝒮* within a distance 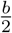 of 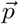 in dimension *j* in the every-epitope model.

Then, the any-epitope model: Let 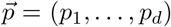 be a reference point in antigenic space. Suppose the strain 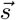 is in *𝒮*, and for all *j*, we have 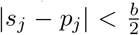. Suppose the host is infected by (or vaccinated with) strain 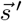 that is also within a distance 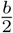 from 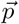 in all dimensions *j* = 1, …, *d*. Applying the same argument as for the every epitope model to each dimension separately, it follows that 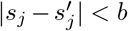 for all *j*, and thus 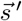 is not added to *𝒮*. Thus 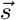 remains the only strain in *𝒮* that is within a distance of 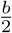 of 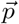 in every dimension in the any-epitope model.

### Immune blind spots calculations

We derive the existence of immune blind spots in the large- 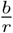 limit, and extend the argument to explain how Fig. 5C and D were generated. For both the every- and any-epitope model, immune blind spots exist because the every- and any-epitope conditions prevent memories from being too closely spaced. The arguments thus proceed by determining, given a strain 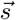 in the memory profile, the maximum possible protection that could be achieved at specially chosen points 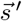 in the case of a maximally “dense” memory profile near 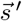.

In the every-epitope model, if strain 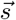 is in a host’s memory profile *𝒮*, no strain 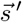 that differs from 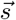 by a distance less than *b* in at least one dimension can be in the memory profile. Let 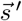 be a strain with 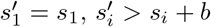 for all *i* = 2, …, *d*, and (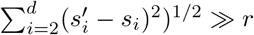, as shown in Fig. 5A in the case where 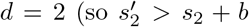 and 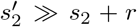). Note that the host with memory (only) at 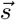 is essentially not protected at 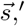, as its Euclidean distance from 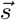 is much greater than the cross-protection distance, and the strain 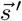 cannot be added to the memory profile *𝒮* even if the host is infected by 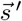, since 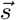 and 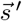 share the first epitope. The closest strains to 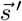 that could be added to *𝒮* are the two strains displaced from 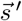 by *b* in the first epitope (with coordinates 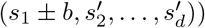, and the strains are mutually exclusive; that is, at most one of these strains can be added without violating the every-epitope condition in epitopes 2 through *d*. If the blunting distance *b* is sufficiently large compared to the cross-protection distance *r*, the maximum protection the individual could have at 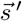 is well-approximated by the protection conferred by one of these two closest strains, even if the memory profile *𝒮* also contains strains further away. The minimum probability of infection upon exposure to 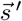 is thus approximately 1 − *e*^*−b/r*^, which approaches 1 as *b*/*r* becomes large. The host thus has an immune blind spot at 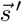 (Fig. 5A) in the large-*b*/*r* limit.

A similar observation holds for the any-epitope model. An individual with strain 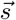 in their memory profile *𝒮* cannot have specific memory to any strain 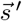 whose coordinates are all within a distance *b* of 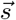. Let 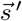 be the strain with coordinates 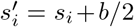 for *i* = 1, …, *d*. In addition to 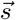, there are (2^*d*^ −1) strains that differ from 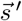 by a distance of *b*/2 in each epitope: these are strains that are located at either *s*_*j*_ or *s*_*j*_ + *b* in each direction *j* (with at least one coordinate differing from 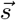). Each of these strains is distance *d*^1*/*2^*b*/2 from 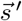, and these are the closest strains to 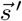 that could be in *𝒮*. As in the every-epitope case, for sufficiently large *b*/*r*, the maximum protection the individual could have at 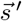 is well-approximated by the protection conferred by these closest strains if memory had been acquired to all of them. The minimum probability that an individual with 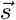 in their memory profile is infected by 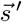 upon exposure therefore equals the probability that neither 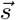 nor any of the other closest strains is protective, which is equal to 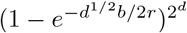. This is close to 1 when *b*/*r* is very large, meaning that the host has an immune blind spot at 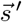 (Fig. 5B).

In reality, we don’t expect the blunting distance to be many orders of magnitude greater than the cross-protection distance. If the blunting distance is not too much larger than the cross-protection distance, so that the large-*b*/*r* limit does not hold, the immune blind spots may cease to be blind spots, as cross-protection from allowed strains (outside the 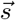 excluded region) can be high enough to provide protection at 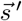. In this case, to determine the existence of immune blind spots, it may not be enough to compute only the protection conferred from the closest allowed strains, as the contribution to protection at 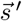 from further allowed strains may be nonnegligible. We therefore consider how much protection from further allowed strains can contribute at 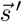 in both the every- and any-epitope models in order to calculate the maximum achievable protection at the points that become blind spots in the large-*b*/*r* limit considered above.

In the every-epitope model, the coordinates of the nearest allowed memories are all equal to 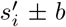; however at most one such memory can be added. The next-nearest memories are distance 2*b* away in the first coordinate, and distance *b* in the remaining (*d*−1) coordinates. In general, for the *k*th-nearest memories, the first coordinate is distance (*kb* + 1) away and the remaining (*d* − 1) coordinates are distance *kb* away, and only one such point can be added for each *k* (without violating the every-epitope condition), no matter the number of dimensions. Each of these points is distance ((*d* − 1) * (*kb*)^2^ + (*kb* + 1)^2^)^1*/*2^ from 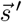, and so the probability it protects against infection at 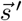 equals 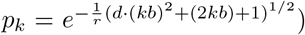. If all of these strains were in the individual’s memory profile, the total probability of infection at 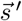 would be equal to the product Π_*k*_(1 − *p*_*k*_), a quantity which is very close to 1 for *b*/*r* ≳ 5 (*d* = 2), 2 (*d* = 3), 1.8 (*d* = 4), etc. (Fig. 5C).

In the any-epitope model, blind spots occur in the center of a *d*-dimensional hypercube, and the densest memory possible profile containing a strain at the origin has strains at each point on a lattice with edges of distance *b*. The probability of protection given the densest possible distribution of memory is thus the product over lattice points of 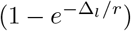, where Δ_*l*_ is the distance from the point 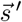 to the lattice point *l*. To compute the probability of infection plotted in Fig. 5D, we explicitly computed the given product over all lattice points up to a distance sufficiently far from the immune blind spot such that adding further lattice points does not appreciably impact the probability of protection. Note that the number of such lattice points at each distance depends on the number of epitopes *d* in a complicated way, and so we calculated separately for each *d*. The associated minimum probability of infection at the blind spot for an individual with memory at 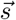, computed by subtracting the probability of protection from 1, is shown in Fig. 5D.

### Details of transmission bound calculation

The general strategy for determining the transmission bound for either blunting model is to impose the OAS constraint over blunting neighborhoods of the appropriate geometry, find the neighborhood over which the integral of the memory distribution is maximized, carry out the integral, and determine the consistency conditions for the inequality to be satisfied.

In the every-epitope model, the *i*th blunting neighborhood about a point 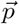 takes the form 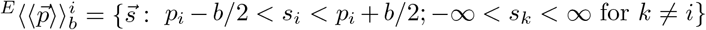. The memory distribution in the successive strain regime falls off exponentially into the past from the wavefront *x*_1_ = *vt*. While blunting neighborhoods are infinitely extended rectangles parallel to the axes, the direction in which the wave travels is not generally parallel to the axes, so the amount of memory captured in the blunting neighborhoods depends on the direction the wave is traveling. The blunting neighborhood over which the integral of memory is maximized is parallel to the axis of the slowest-evolving epitope, with the wavefront at one of its corners (Fig. 8A,B). The memory captured therefore corresponds to the interval 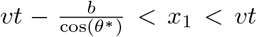 in wave-adapted coordinates. The every-epitope OAS constraint simplifies to

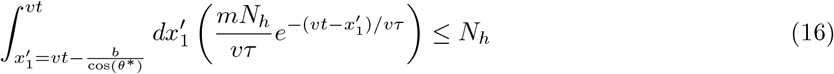

In the any-epitope model, the blunting neighborhood about a point 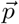 takes the form 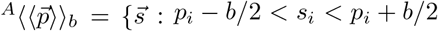 for *i* = 1, …, *d*}. These blunting neighborhoods are *d*-dimensional cubes with side length *b* and sides parallel to the axes. As in the every-epitope model, the amount of memory captured in the blunting neighborhoods depends on the direction the wave is traveling. For the any-epitope geometry, the blunting neighborhood over which the integral of memory is maximized again has the wavefront at one of its corners. This time, however, the length of the interval captured in the neighborhood depends on the direction of the fastest-evolving epitope. The memory captured still corresponds to the interval 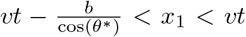 in wave-adapted coordinates (Fig. 8C,D), but now *θ*^*^ indicates the fastest-evolving rather than the slowest-evolving direction. The any-epitope OAS constraint simplifies to

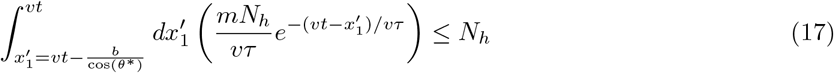

Both of these integrals are of the form 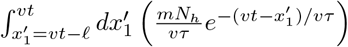 with different values of *𝓁*, so we evaluate this integral and then substitute for *𝓁* to obtain the relevant inequalities for each blunting model. The inequality simplifies to the condition

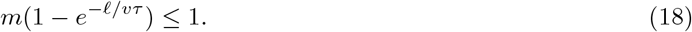

This is equivalent to the condition that

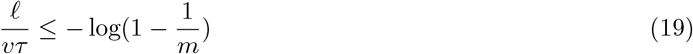

The derived quantity *vτ* can be expressed in terms of the fundamental parameters that characterize the system by manipulating several expressions derived in [36]. The timescale *τ* equals *mN*_*h*_/*N*_*I*_. The incremental fitness gradient along the direction of motion of the wave is analogous to the fitness effect of mutations, *s*, and equals 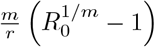 in the traveling wave system. Marchi et al. argue that *s* is related to *v* via the relation 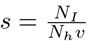. It follows that

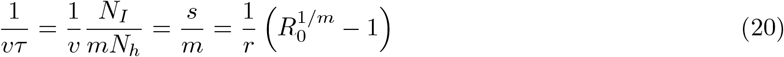

so that the inequality in Eq. 19 becomes

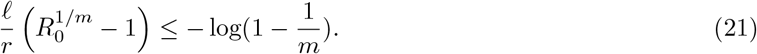

The limit as *m* becomes large of 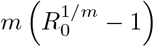 is equal to log *R*_0_, and more generally the following inequality holds for any value of *m* greater than 1,

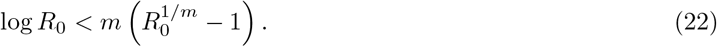

Together, Eq. 21 and Eq. 22 imply that the following inequality among the fundamental parameters that characterize transmission and the immune response must hold,

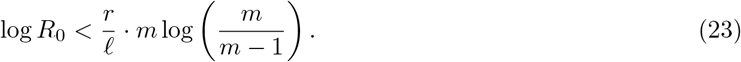

In the large-*m* limit, Eq. 23 simplifies to

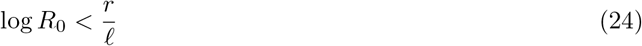

This is the same as the expressions given in the Results, where *𝓁* equals 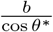.

**Figure 8.**
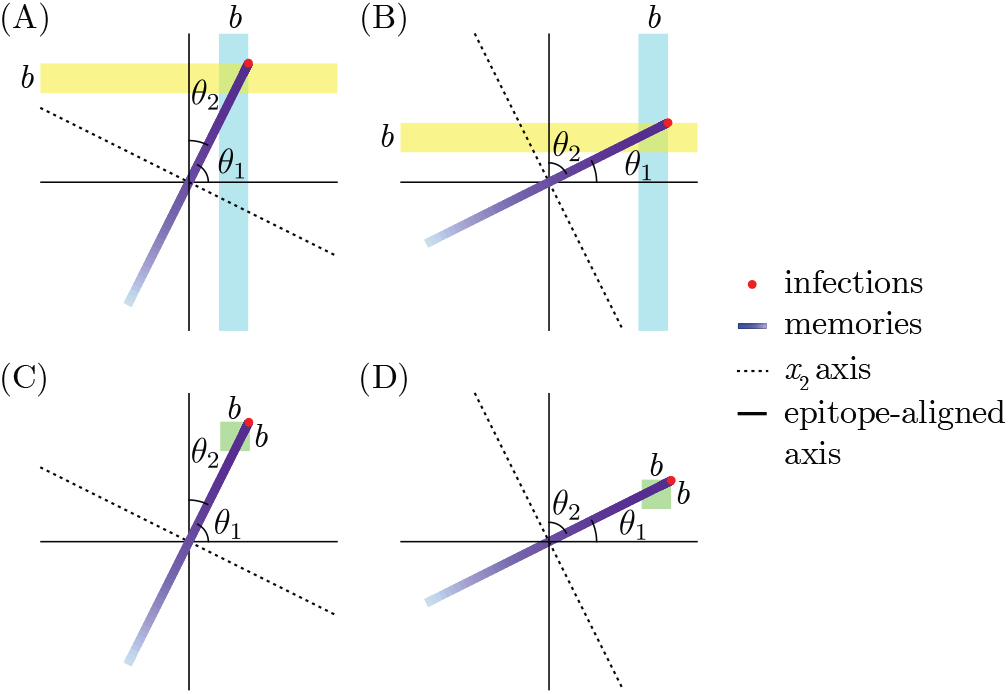
The slowest (fastest) epitope determines the integration bounds in the every- (any-) epitope model. (A, B) The every-epitope model. (A) The blue stripe is the blunting neighborhood that maximizes the number of memories being integrated over. The interval of *x*_1_ that lies in the blue region has length *b*/ cos *θ*_1_. The projection of the velocity vector onto the epitope 1 direction is less than the projection onto the epitope 2 direction, confirming that *θ*_1_ is the angle corresponding to the slowest evolving epitope. (B) The yellow stripe is the blunting neighborhood that maximizes the number of memories being integrated over. The interval of *x*_1_ that lies in the yellow region has length *b*/ cos *θ*_2_. (C, D) The any-epitope model. The green regions are the blunting neighborhoods that maximize the number of memories being integrated over. The length of the *x*_1_ interval that lies in the green region equals *b*/ cos *θ*_2_ in (C), and *b*/ cos *θ*_1_ in (D).

### Bounds on dynamical quantities in traveling wave model

In this section, we give the expressions for dynamical quantities in the antigenic space model, originally derived in [36], and numerically demonstrate the monotonicity of these quantities in the ratio 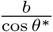, implying *D*-dependent extrema when *R*_0_ equals the limiting value 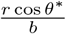, as in Eq. 14.

The quantities *v, σ* and *t*_MRCA_ are simply expressed in terms of the fitness increment 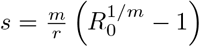:

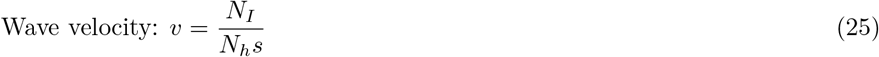

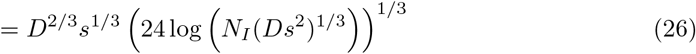

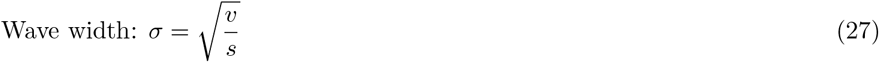

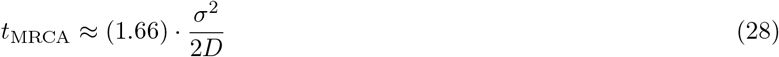

Note that the wave velocity is defined implicitly through Eq. 25 and Eq. 26, which determine both *v* and the number of infected hosts *N*_*I*_ in terms of the independent parameters of the system (*m, R*_0_, *R, D, N*_*h*_).

The quantities *v* (Fig. 9A), *σ* (Fig. 9B), and *t*_MRCA_ (Fig. 92C) are monotonic in the fitness gradient *s*. The bound on *R*_0_ for serial strain replacement, Eq. 14, is equivalent to the statement that 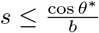 in the large- *m* limit, since 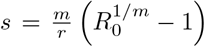 approaches 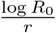 and 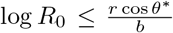. It follows that a monotonically increasing (decreasing) function of *s* is maximized (minimized) over all traveling wave solutions at the limiting value 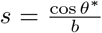. Thus for any values of *D, b, θ*^*^, and *N*_h_, there exists a maximum limit on *v* and minimum limits on *σ* and *t*_MRCA_ which are independent of *m* in the large-*m* limit. These bounds are plotted in the left panels of Fig. 7.

**Figure 9.**
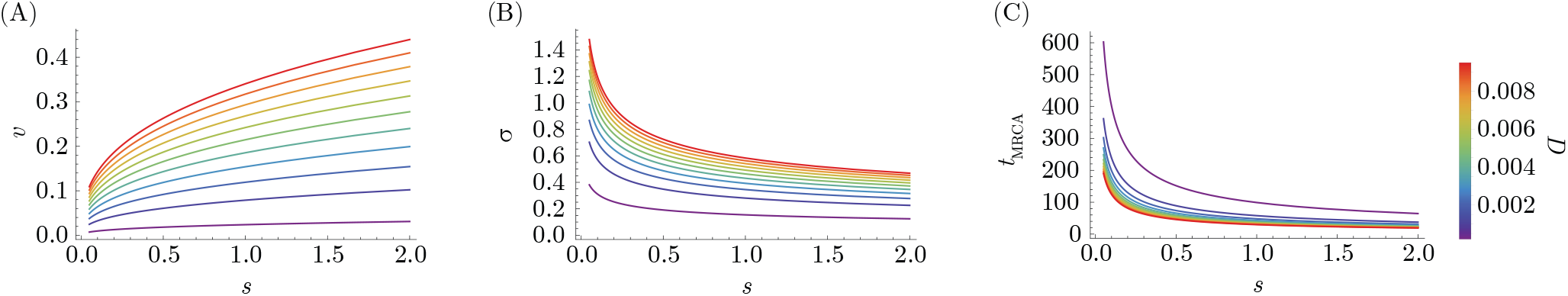
Dynamical quantities are monotonic as a function of the fitness gradient *s*. A) The wave velocity is monotonically increasing, and (B, C) the infection standard deviation *σ* as well as the time to most recent common ancestor *t*_MRCA_ are monotonically decreasing.

## Code Availability

All code used for analytical calculations and generation of figures will be available upon publication. The code is on GitHub at https://github.com/unrealmcg/OAS-strain-replacement/tree/main.

## Acknowledgments

We thank Manon Ragonnet, Christopher Joel Russo, Maryn Carson, and Tiffany Cai for useful discussions and feedback on the manuscript, and Christopher Joel Russo for help with formatting figures. Research reported in this publication was supported by the National Institute of the General Medical Sciences and the National Institute of Allergy and Infectious Diseases of the National Institutes of Health under award numbers F32GM134721 and R01AI149747. The content is solely the responsibility of the authors and does not necessarily represent the official views of the National Institutes of Health. SC acknowledges support from the National Institute for Theory and Mathematics in Biology through the National Science Foundation (grant number DMS-2235451) and the Simons Foundation (grant number MP-TMPS-00005320).

## Footnotes

1 See Methods, Eq. 23, for the finite-*m* correction, which quickly approaches the large-*m* behavior.

